# Dynamic analysis of pulsed cisplatin identifies effectors of resistance in lung adenocarcinoma

**DOI:** 10.1101/775924

**Authors:** Jordan F. Hastings, Alvaro Gonzalez-Rajal, Jeremy Z.R. Han, Rachael A. McCloy, Yolande E.I. O’Donnell, Monica Phimmachanh, Alexander D. Murphy, Adnan Nagrial, Dariush Daneshvar, Venessa Chin, D. Neil Watkins, Andrew Burgess, David R. Croucher

## Abstract

Identification of clinically viable strategies for overcoming resistance to platinum chemotherapy in lung adenocarcinoma has been hampered by inappropriately tailored *in vitro* assays of drug response. Therefore, using a pulse model that closely recapitulates the *in vivo* pharmacokinetics of platinum therapy, we profiled cisplatin-induced signalling, DNA damage and apoptotic responses across a panel of lung adenocarcinoma cell lines. By coupling this data with real-time, single cell imaging of cell cycle and apoptosis, we show that *TP53* mutation status influenced the mode of cisplatin induced cell cycle arrest, but could not predict cisplatin sensitivity. In contrast, P70S6K-mediated signalling promoted resistance by increasing p53/p63 and p21 expression, reducing double-stranded DNA breaks and apoptosis. Targeting P70S6K sensitised both *TP53* wildtype and null lines to cisplatin, but not *TP53* mutant lines. In summary, using *in vitro* assays that mimic *in vivo* pharmacokinetics identified P70S6K as a robust mediator of cisplatin resistance and highlighted the importance of considering somatic mutation status when designing patient-specific combination therapies.

## Introduction

Lung adenocarcinoma is the most common form of lung cancer, the leading cause of cancer-related death worldwide. Lung adenocarcinoma is typically diagnosed late, meaning that most patients require systemic chemotherapy (Chen et al., 2014). Platinum-based chemotherapy is likely to remain an important treatment modality for these patients due to the emergence of resistance to targeted therapies in EGFR, ALK or ROS mutant tumours (Lindeman et al., 2018), and the fact that most patients do not respond to single agent immunotherapy (Doroshow et al., 2019, Kim et al., 2019).

Despite the use of platinum-based chemotherapy in lung adenocarcinoma for over four decades, response rates remain below 30% due to the prevalence of innate resistance (Bonanno et al., 2014, Pilkington et al., 2015). In addition, dose-related nephrotoxicity remains a challenge in many patients (Pabla & Dong, 2008). Strategies to improve platinum efficacy could therefore significantly improve outcomes for lung adenocarcinoma patients. However, unravelling platinum resistance in lung adenocarcinoma has proven challenging, as over 147 mechanisms of resistance have been proposed (Stewart, 2007), yet there remains a lack of viable clinical options to improve response rates.

From an experimental viewpoint, discordance between the *in vivo* pharmacokinetics of platinum chemotherapies and their use within *in vitro* assays has likely contributed to the identification of putative resistance mechanisms and drug targets that have not ultimately translated to the clinic. We have previously shown that predictive models of drug-induced apoptotic signalling dynamics can be used to stratify cancer patients (Fey et al., 2015). To apply a similar concept to platinum resistance, we first considered that traditional *in vitro* methods have involved measures of drug response in cancer cells cultured in the continuous presence of high-dose chemotherapy over several days. This contrasts with pharmacokinetic studies in humans and rodents demonstrating that both cisplatin and carboplatin are rapidly cleared from the circulation, and the tumour, within 2-3 hours following administration (Andersson et al., 1996, Johansen et al., 2002). Therefore, in this study we have utilised a cisplatin pulse model which more closely recapitulates these physiological pharmacokinetics, aiming to maintain the fidelity of the apoptotic mechanism mediated by cisplatin *in vivo*.

We now present an in-depth analysis of the dynamic signalling response to a pulse of platinum chemotherapy, describing the relationship between a number of key signalling nodes, the DNA damage response and platinum sensitivity. Importantly, we also propose a therapeutic strategy targeting P70S6K using the dual PI3K/mTOR inhibitor dactolisib, with the potential to improve the efficacy of current platinum based treatment regimens.

## Results

### Continuous versus pulsed cisplatin treatment

In order to directly compare the response of lung adenocarcinoma cells to the continuous presence of cisplatin, or a pulse of cisplatin that mimics *in vivo* pharmacokinetics (2h, 5 μg/mL) (Figure 1A), we monitored the growth and apoptosis of the innately resistant A549 lung adenocarcinoma cell line (Marini et al., 2018) by live cell imaging under both conditions (Figure 1B). This analysis demonstrated that while continuous exposure to cisplatin resulted in decreased cell number and increased apoptosis over 72h, a pulse of cisplatin only reduced the rate of cell proliferation and did not induce apoptosis.

**Figure 1:**
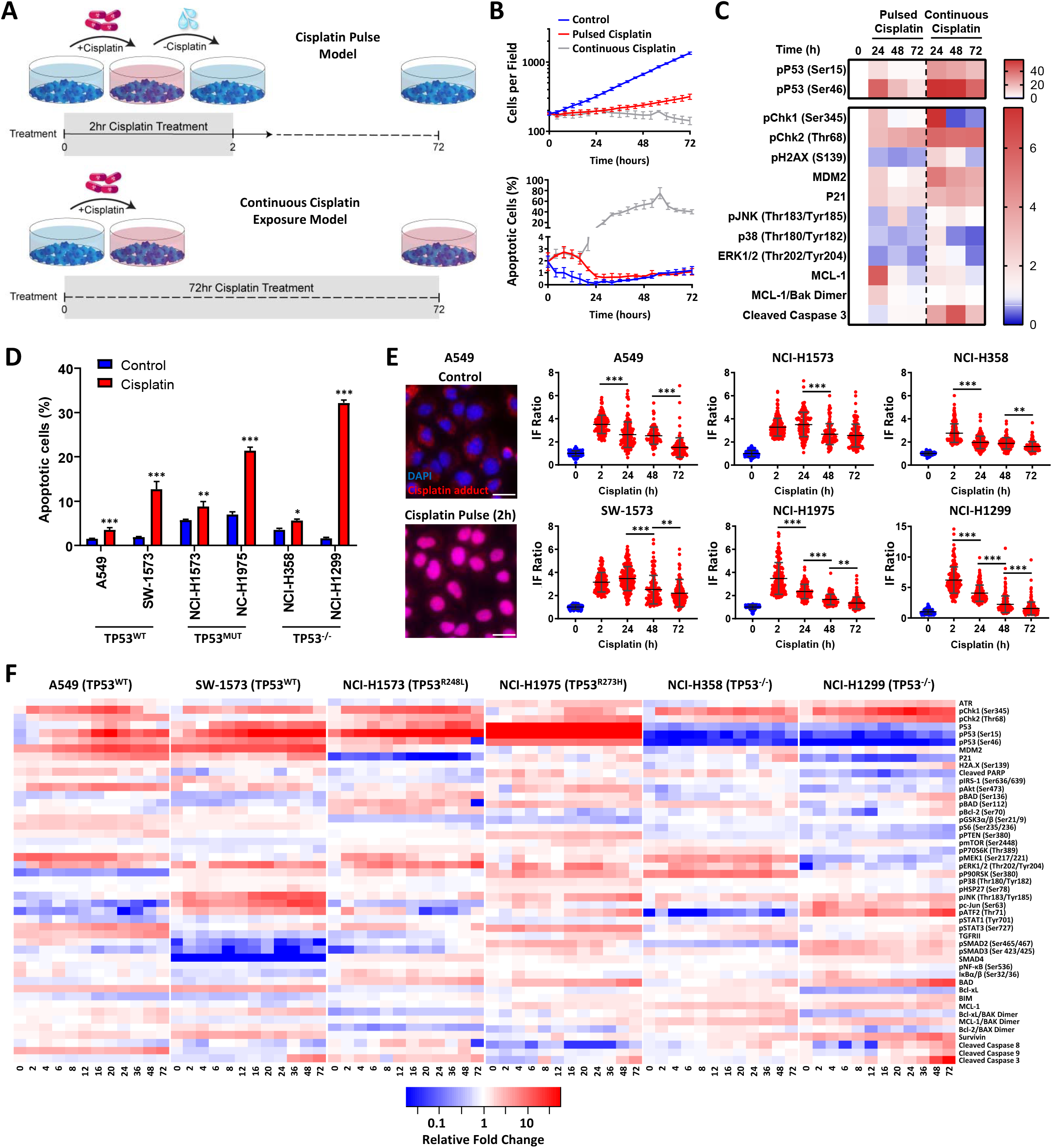
Multiplexed analysis of cisplatin-induced signalling. **(A)** Schematic of the cisplatin pulse model (5 µg/mL, 2 h) and continuous pulse model (5 μg/mL, 72 h). **(B)** Live-cell imaging of A549 cells treated with either continuously, or with a cisplatin pulse. Apoptotic cells were identified by uptake of propidium iodide (mean ± SD). **(C)** Multiplexed analysis of key DNA damage, apoptosis and signalling proteins in A549 cells treated either continuously, or with a cisplatin pulse (n=3, mean). **(D)** Apoptosis measured by propidium iodide staining for the sub-G1 population, performed 72 h following a cisplatin pulse across a panel of lung adenocarcinoma cell lines, as indicated (n=3, mean ± SD, *** p<0.001, ** p<0.01, * p<0.05). **(E)** Representative images of anti-cisplatin antibody staining in A549 cells following a cisplatin pulse, and quantification of nuclear cisplatin-DNA adducts across the cell line panel (n≥100, mean ± SD. *** p<0.001, ** p<0.01). All treatment conditions (red) are significantly different from control (blue), p<0.001. **(F)** Multiplexed analysis of DNA damage, apoptosis and signalling pathways following a cisplatin pulse across a panel of lung adenocarcinoma cell lines, as indicated (n=3, mean).

To further examine the differences between these two models, we used multiplexed, bead-based protein analysis to investigate the DNA damage, apoptotic and signalling response for key pathway components previously implicated in the response to continuous cisplatin exposure (I.e. p38, ERK, JNK and MCL-1) (Marini et al., 2018, Stewart, 2007) (Figure 1C, Supplementary Figure 1). As might be expected, the continuous exposure model resulted in a significantly elevated and sustained DNA damage response when compared to the pulse model, particularly for the phosphorylation of Chk2 (Ser345), p53 (Ser15 and Ser46), pH2A.X (Ser139 - γH2A.X) and expression of p21 and MDM2. This heightened DNA damage response during the continuous exposure to cisplatin was also reflected in the increased activation of Caspase 3, which was completely absent for the pulse model. Furthermore, while p38 and ERK activation were significantly increased in cells continuously exposed to cisplatin, the expression of MCL-1 and detection of MCL-1/Bak dimers only significantly increased in cells treated with a pulse of cisplatin. This finding demonstrates that not only does the continuous exposure model result in a DNA damage and apoptotic response that is incongruent with that observed following a pulse of cisplatin, the dynamics of key signalling pathways are fundamentally different between these two treatment models. Taken together, this data demonstrates that previous mechanisms of platinum resistance established using a continuous exposure model may be based on artefacts arising from non-physiological levels of drug exposure.

### Response to cisplatin is not associated with either TP53 status or drug-efflux

To investigate potential mechanisms of platinum resistance using a model consistent with the physiological pharmacokinetics of platinum therapy, we applied this cisplatin pulse model to a panel of six lung adenocarcinoma cell lines with distinct *TP53* mutation backgrounds (two wildtype lines, two mutant *TP53* lines and two *TP53* null) and measured the apoptotic response at 72 h (Figure 1D). Based upon this model we observed a range of sensitivity to cisplatin, from the most resistant A549 line (~3% apoptosis) to the most responsive NCI-H1299 line (~32% apoptosis). However, these cell lines could not be stratified simply according to their *TP53* mutation status, or other frequently observed genetic alterations (Supplementary Table 1).

As the action of drug-efflux pumps is another commonly proposed mechanism of resistance to platinum therapy (Hoffmann & Lambert, 2014), we performed fluorescence microscopy with an antibody towards cisplatin-induced DNA adducts at multiple time-points following a 2 h cisplatin pulse (Figure 1D). This analysis demonstrated that within this model, all six cell lines displayed significant nuclear localised cisplatin-DNA adducts following a 2 h pulse (Figure 1E, Supplementary Figure 2), suggesting that drug efflux is not associated with variations in the apoptotic response to a pulse of cisplatin in these lines. Furthermore, these cisplatin-DNA adducts progressively resolved over a 72 h period in all cell lines (Figure 1C), confirming that pathways responsible for facilitating the removal of cisplatin adducts are also functional across this panel.

**Figure 2:**
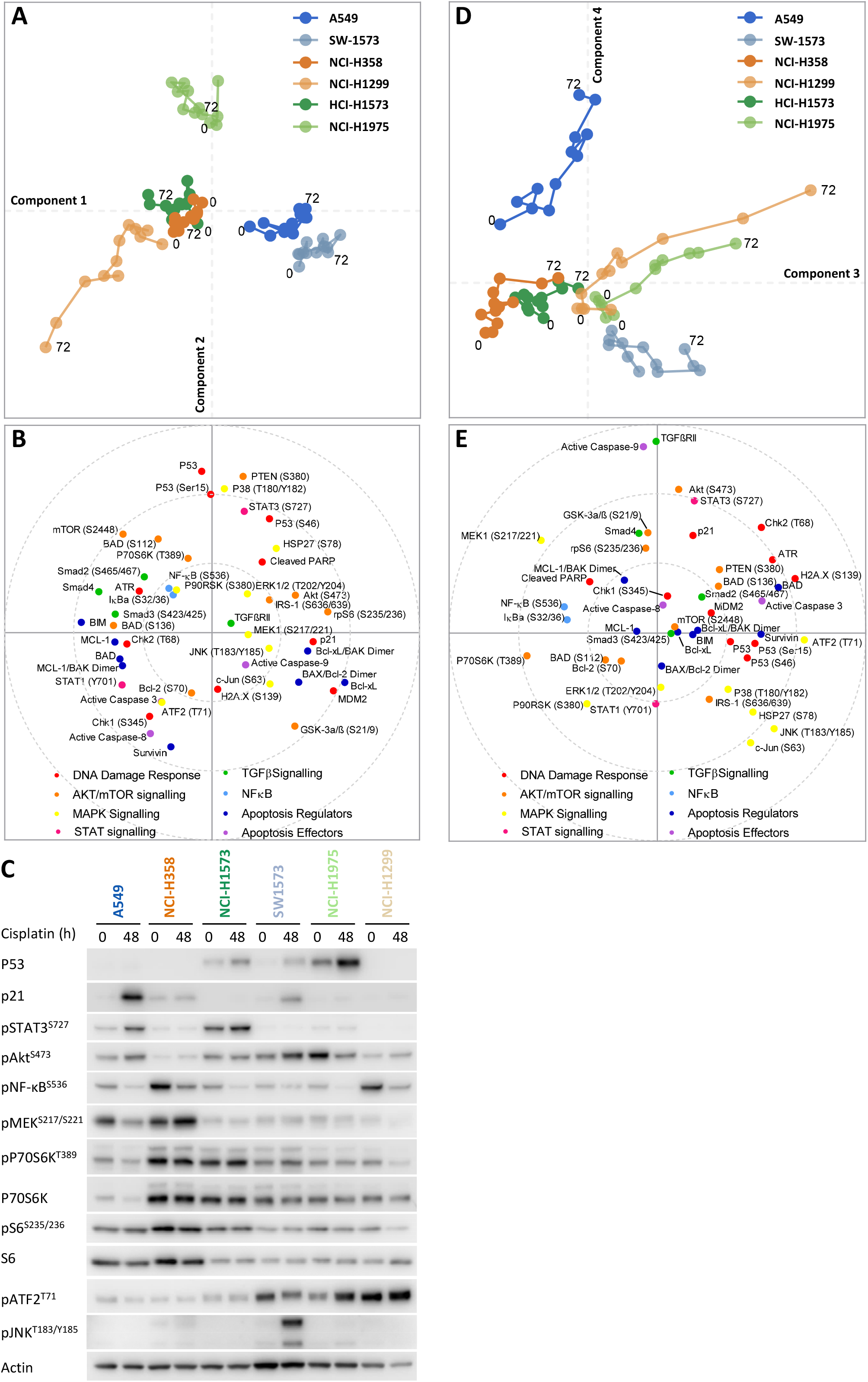
Principal component analysis. **(A)** Visualisation of component 1 against component 2 for the cisplatin induced signalling analysis across the cell line panel. **(B)** Distribution of the analytes according to their weighting within component 1 and 2. **(C)** Western blotting for selected analytes across the cell panel prior to, and 48 h post a cisplatin pulse. **(D)** Visualisation of component 3 against component 3 for the cisplatin induced signalling analysis across the cell line panel. **(E)** Distribution of the analytes according to their weighting within component 1 and 2.

### Multi-dimensional analysis of cisplatin-induced signalling dynamics

To gain an understanding of the wider signalling networks associated with resistance to platinum therapy, we utilised multiplexed, magnetic bead based assays to profile the signalling and DNA damage response at multiple time points following a 2 h pulse of cisplatin, across all six cell lines (Figure 1F). For this analysis, we tracked the dynamics of 47 different protein analytes (Supplementary Table 2) over a 72 hour period, focusing on elements of signalling network structures that we recently implicated in platinum chemoresistance (Marini et al., 2018), including the MAPK, PI3K/mTOR, NF-κB and TGFβ pathways, as well as a number of key apoptotic mediators and DNA damage response proteins (Figure 1G).

From this dataset, a clear correlation can be seen between the *TP53* mutation status of each cell line and the dynamics of the p53 pathway (Supplementary Figure 3), which validates the fidelity of the multiplexed analysis platform for this form of analysis. In line with the detection of cisplatin-DNA adducts (Figure 1E), Chk1 (Ser345) phosphorylation increased rapidly in all lines following the 2 h pulse, followed by a slower wave of Chk2 (Thr68) phosphorylation in all lines except SW1573. Within both the *TP53* wildtype cell lines (A549 and SW-1573) this was followed by phosphorylation of p53 (Ser15 and Ser46), the accumulation of total p53 and increased expression of the p53 transcriptional targets MDM2 and p21. In the *TP53* mutant cell lines (NCI-H1573 and NCI-H1975) p53 phosphorylation still occurred, however in line with their loss of DNA binding capability, this did not result in increased expression of MDM2 or p21. As would be expected for the *TP53* null lines (NCI-H358 and NCI-H1299), p53 is absent and therefore not detected in either the total or phosphorylated form. Interestingly though, there was a significant increase in p21 and MDM2 expression in the NCI-H358 line in the absence of p53 expression.

**Figure 3:**
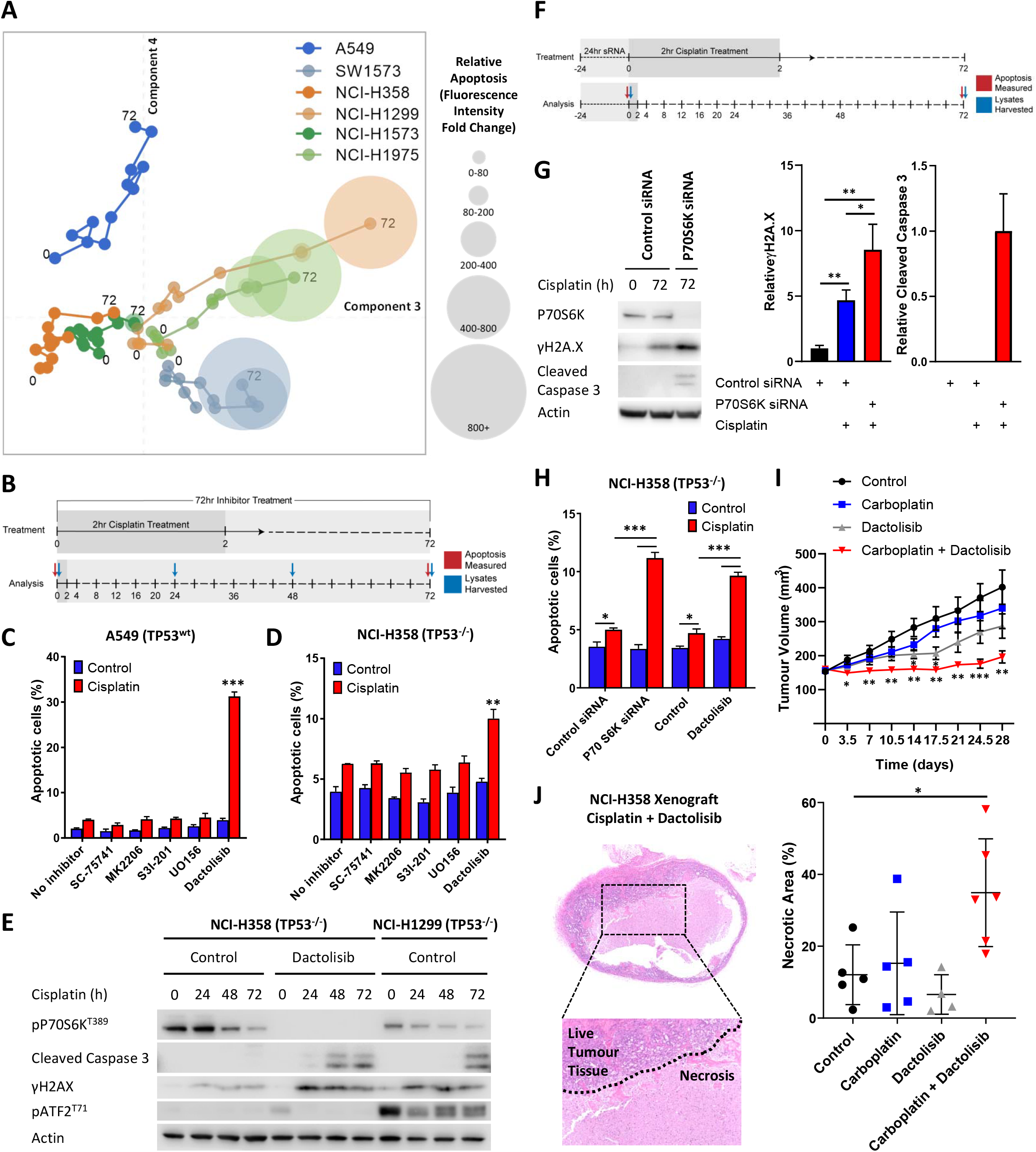
Visualising and targeting mechanisms of cisplatin resistance. **(A)** An overlay of live-cell imaging of apoptosis following a cisplatin pulse on top of the PCA plot of component 3 against component 4. **(B)** Schematic of the cisplatin pulse model, with the addition of small molecule inhibitors, and outline of sample collection for apoptosis assays and western blotting. **(C,D)** Apoptosis measured by propidium iodide staining for the sub-G1 population in A549 and NCI-H358 cells, performed 72 h following a cisplatin pulse with the addition of small molecule inhibitors (1 µM) as indicated (n=3, mean ± SD). **(E)** Western blotting on lysates from NCI-H358 and NCI-H1299 cells following a cisplatin pulse, with the addition of dactolisib (1 µM), as indicated. **(F)** Schematic of the cisplatin pulse model, with the addition of siRNA pre-treatment, and outline of sample collection for apoptosis assays and western blotting. **(G)** Western blotting on lysates from NCI-H358 cells, treated with P70S6K or control siRNA, as indicated, prior to and following a cisplatin pulse (n=3, mean ± SD). **(H)** Apoptosis measured by propidium iodide staining for the sub-G1 population in NCI-H358 cells, treated with P70S6K, control siRNA or dactolisib, as indicated, performed 72 h following a cisplatin pulse (n=3, mean ± SD). **(I)** Tumour growth in nude mice bearing NCI-H358 xenografts with continuous treatment of vehicle control or dactolisib (45 mg/kg) prior to, and following a single dose of carboplatin (60 mg/kg) (n≥4, mean ± SEM). **(J)** Quantification of necrosis in NCI-H358 xenografts following the treatment described in (I) (n≥4, mean ± SD). For all experiments, *** p<0.001, ** p<0.01, * p<0.05.

As we had already determined that *TP53* status alone was not sufficient to explain resistance to cisplatin (Figure 1D), we further analysed the whole dataset by performing a principal component analysis (PCA) (Figure 2). This analysis created a visual representation of the dynamic association between key signalling nodes and the response to cisplatin across the entire cell line panel. Using this multidimensional analysis, we were able to capture ~70% of variance in the dataset within the first 4 principal components (Supplementary Figure 4A). Unsurprisingly, plotting the first two principal components against each other (Figure 2A) resulted in separation of the cell lines primarily according to their *TP53* status. As might be expected, within these components the *TP53* wild-type A549 and SW-1573 lines associated with higher p21 and MDM2 expression, while the *TP53* mutant lines separated from the *TP53* null lines mostly on the basis of higher p53 expression and phosphorylation levels (Figure 2A,B,C). However, plotting the third and fourth principal components (Figure 2D,E) created a clear delineation between the three most resistant cell lines (A549, NCI-H358 and NCI-H1573), which cluster towards the left hand side of component 3 (x-axis), and the three most sensitive lines (SW-1573, NCI-1975 and NCI-H1299) which move progressively along component 3 over the 72 h timeframe.

**Figure 4:**
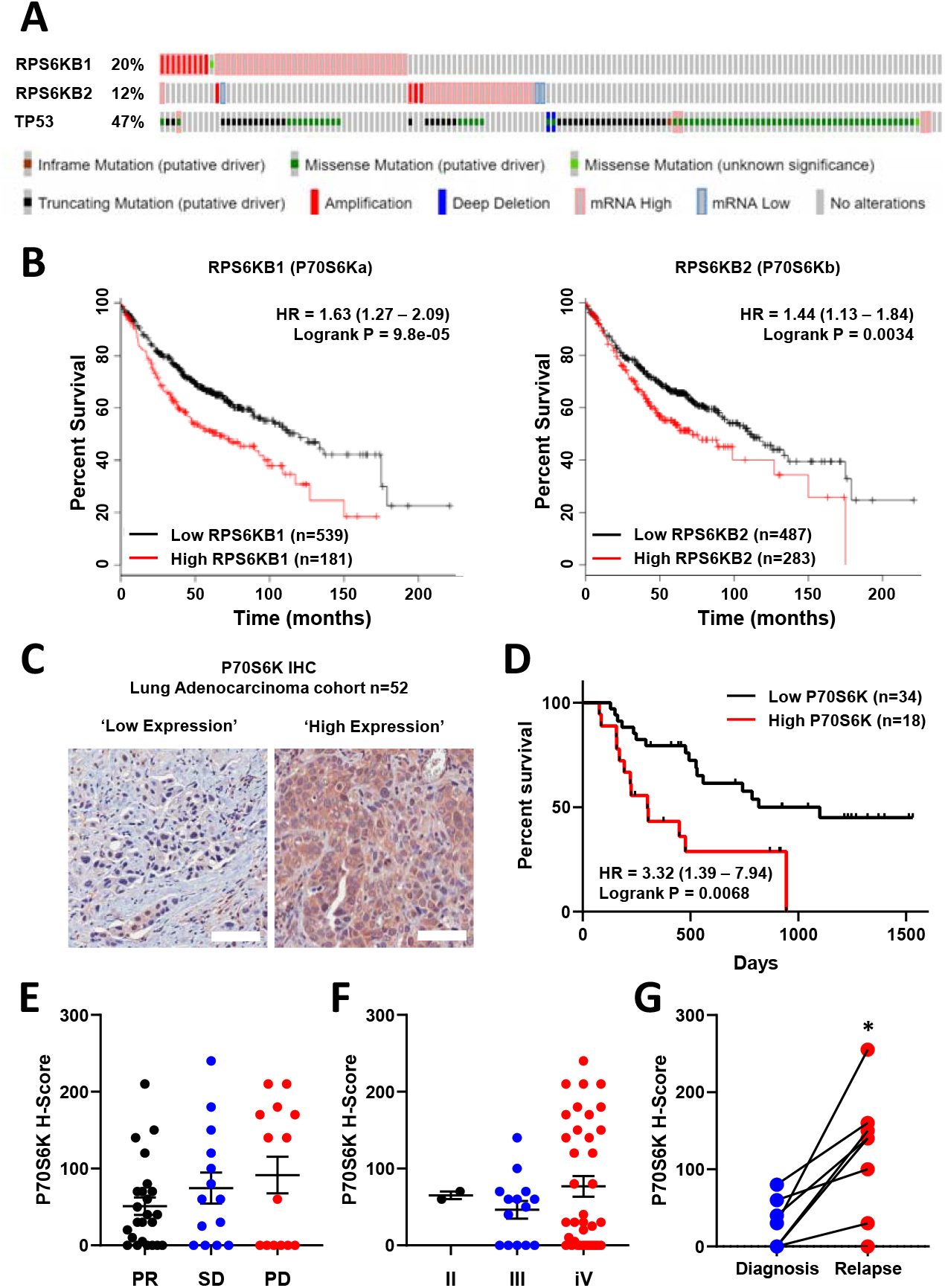
P70S6K in lung adenocarcinoma. **(A)** The frequency of somatic mutations and mRNA over-expression for the *RPS6KB1*, *RPS6KB2* and *TP53* genes in a publically available cohort of lung adenocarcinoma patients (cBioPortal). **(B)** The association between *RPS6KB1* and *RPS6KB2* mRNA expression and overall patient survival in a publically available patient cohort (KM Plotter). **(C)** Representative images of P70S6K immuno-histochemistry staining from a cohort of 52 lung adenocarcinoma patients (Scale bar = 100 µM). **(D))** Survival analysis based upon P70S6K staining in this cohort. **(E)** The association between tumour P70S6K staining intensity (H-Score) and patients that underwent a partial response (PR), or presented with stable disease (SD) or progressive disease (PD). **(F)** The association between tumour P70S6K staining intensity and tumour stage. **(G)** P70S6K staining in 8 matched patient samples from diagnosis and relapse (n=8, * p<0.05).

### Validation of model-based observations

As platinum chemotherapies work by forming covalent DNA adducts, which distort the DNA helix and block replication, the progressive accumulation of single stranded and double stranded breaks is thought to induce apoptosis (Jamieson & Lippard, 1999). This is in line with the movement of cisplatin sensitive cell lines along component 3 (x-axis) towards higher levels of γH2A.X (H2A.X^S139^), cleaved caspase 3 and a stress associated MAPK signalling axis (pATF2, pJNK, pc-Jun) (Figure 2D,E). The association between this signalling state and increased sensitivity to cisplatin can also be clearly observed by overlaying an orthogonal readout of the apoptotic response onto this PCA plot (Figure 3A), generated using live cell imaging to measure caspase activity as an indicator of cell death for 72 h following the cisplatin pulse treatment.

Using this approach, we now created a visual representation that both reflects the variance within the original dataset and demonstrates the key signalling nodes that are associated with differing degrees of platinum-induced apoptosis. While increasing levels of apoptosis are observed over time in the three sensitive cell lines (SW-1573, NCI-1975 and NCI-H1299), the three resistant cell lines display significantly lower levels of apoptosis (Supplementary Figure 4B) and do not move along component 3 towards the region of DNA damage and apoptosis. Instead the NCI-H358 and NCI-1573 lines remain towards the left hand side of component 3, in a region characterised by higher phosphorylation of the mTOR pathway component P70S6K (Thr389), the MAPK components MEK1 (Ser217/221) and P90RSK (S380), NF-κB (S536) and IκBα (Ser32/36) (Figure 2D,E). The resistant A549 line also remains shifted towards the left of component 3, although also moves up component 4 towards a region with higher expression of TGFβRII, cleaved caspase 9, pAkt (Ser473) and pSTAT3 (Ser727) (Figure 2D,E). The association of elevated pMEK (Ser217/221), pNF-κB (S536), pSTAT3 (Ser727) and pP70S6K (Thr389) within these resistant cell lines was also further confirmed by western blotting of independent samples (Figure 2C).

The clear separation of resistant and sensitive cell lines within this multi-dimensional analysis suggests an antagonistic relationship between apoptosis promoted through the progressive accumulation of DNA-damage following a cisplatin pulse, and elevated mTOR, MAPK, Akt, STAT or NF-κB signalling events. However, as a statistical process, relationships derived from a principal component analysis are purely correlative in nature and require a further degree of validation before any causative conclusions may be drawn. We therefore sought to identify effectors of platinum resistance by including specific inhibitors of these signalling pathway components during and after the 2 h cisplatin pulse, followed by a measurement of apoptosis at 72 h (Figure 3B). Using the resistant A549 (Figure 3C) and NCI-H358 (Figure 3D) cell lines we observed that specific inhibitors of Akt (MK2206), STAT3 (S3I-201), MEK (UO126) or NF-κB (SC-75741) did not significantly increase apoptosis in either cell line, suggesting that while elevated activity of these signalling proteins may be present in one or both of these resistant lines (Figure 2C), they are not causally associated with resistance to cisplatin. Instead, under these conditions, only the inhibition of P70S6K with the dual PI3K/mTOR inhibitor dactolisib resulted in a significant increase in cisplatin-induced apoptosis in both cell lines. P70S6K is a serine/threonine-specific protein kinase known to require phosphorylation by both PI3K and mTOR for activation (Sunami et al., 2010) (Moser et al., 1997). While dactolisib will result in the inhibition of a number of substrates downstream of both PI3K and mTOR, the lack of sensitisation by an Akt inhibitor (MK2206) suggests a specific role for P70S6K in mediating resistance to cisplatin.

Taking the opposite approach, elevated levels of JNK pathway activity (pJNK, pc-Jun) were also observed following cisplatin treatment within the SW-1573 line (Figure 2C,D,E). However, the inclusion of a JNK inhibitor (JNK inhibitor VIII) did not prevent cisplatin induced apoptosis in this cell line (Supplementary Figure 4C), suggesting that JNK activity was not promoting apoptosis in this context.

### P70S6K promotes platinum resistance in lung adenocarcinoma

Comparing the relative expression levels of phosphorylated P70S6K across our stratified panel of cell lines, higher expression was observed in the resistant NCI-H358 and NCI-H1573 lines (Figure 2C). This expression pattern was also mirrored by the levels of total P70S6K (Figure 2C), which seems to be primarily driving the observed levels of P70S6K phosphorylation. Interestingly, the resistant A549 line did not have elevated P70S6K expression or phosphorylation, although as reflected in the PCA plot (Figure 2E) this cell line did have high expression of the P70S6K substrate, Ribosomal Protein S6, leading to a level of phosphorylation equivalent to that observed in the resistant NCI-H358 line (Figure 2C).

Utilising the highest expressing (NCI-H358) and lowest expressing (NCI-H1299) lines, which are also both *TP53* null, we observed that cisplatin treatment resulted in greater caspase 3 cleavage and γH2A.X expression in the sensitive NCI-H1299 line (Figure 3E), which is in line with the multiplexed signalling analysis (Figure 1G). Crucially, treatment of the resistant NCI-H358 line with dactolisib during and after the cisplatin pulse increased both caspase 3 cleavage and γH2A.X expression to that observed in the sensitive NCI-H1299 line. Conceptually, this now mimics the movement of the resistant NCI-H358 line along component 3 of our PCA analysis, towards the region of DNA damage and apoptosis characterised by the sensitive cell lines (Figure 2, 3A). This finding demonstrates that while this form of multi-dimensional analysis of signalling networks creates a set of correlative relationships, this data can also be utilised to investigate causal effectors of downstream cellular behaviour.

As mentioned above, dactolisib can efficiently inhibit the phosphorylation of P70S6K, but will also result in the inhibition of a number of other potential PI3K/mTOR substrates. To confirm that P70S6K is a key downstream component mediating platinum resistance in this context, we knocked down P70S6K using siRNA in the NCI-H358 line, prior to proceeding with a cisplatin pulse (Figure 3F). Under these conditions, cisplatin treatment resulted in significantly higher caspase 3 cleavage and γH2A.X expression in the P70S6K knockdown cells (Figure 3G). Additionally, P70S6K knockdown also significantly increased cisplatin-induced apoptosis in the NCI-H358 cells, to the same extent as that observed with dactolisib treatment (Figure 3H).

To further validate this combination therapy approach *in vivo* and validate the findings of our pulse model, we treated mice bearing NCI-H358 xenografts with either carboplatin, dactolisib, or a combination of both (Figure 3I). For this model, dactolisib (45 mg/kg) was delivered by oral gavage, prior to a one-off intraperitoneal injection of carboplatin (60 mg/kg), and throughout the course of the experiment. Under these conditions, carboplatin had no significant effect upon tumour growth, while dactolisib had a moderate effect as a single agent that was only significant at days 14 and 17. However the combination of carboplatin and dactolisib completely halted tumour growth, even up to 28 days following the single pulse of carboplatin treatment. Furthermore, an analysis of tumour sections revealed that the carboplatin and dactolisib treated xenografts were not only smaller in size, but also had significantly larger regions of necrosis (Figure 3J).

These findings demonstrate that elevated P70S6K activity can specifically mediate resistance to platinum therapy in lung adenocarcinoma, which can be effectively targeted with the dual PI3K/mTOR inhibitor dactolisib. Importantly, elevated P70S6K activity is frequently observed in several cancer subtypes, including lung cancer (Chen et al., 2017), and its overexpression is commonly associated with aggressive malignant phenotypes and poor overall prognoses (Ip & Wong, 2012). Indeed, an analysis of TCGA data using cBioPortal (Cerami et al., 2012, Gao et al., 2013) reveals that *RPS6KB1* and *RPS6KB2*, the two isoforms of P70S6K, are amplified or over-expressed in 20% and 11% of lung adenocarcinoma cases, respectively (Figure 4A). Importantly, while *TP53* mutations or deletions also occur within 47% of lung adenocarcinomas, *RPS6KB1* and *RPS6KB2* amplification/over-expression occur on the background of either wildtype or mutant *TP53*.

Further analysis with KM plotter (Gyorffy et al., 2013) demonstrated that the elevated mRNA expression of both isoforms was also significantly associated with poor overall survival of lung adenocarcinoma patients (Figure 4B). We therefore investigated the association with survival and response to chemotherapy by performing IHC for total P70S6K in a cohort of 52 lung adenocarcinoma patients that all received a neoadjuvant chemotherapy regimen containing platinum-based chemotherapy (Figure 4C, Supplementary Table 3). In this cohort, high expression of P70S6K was significantly associated with poor overall survival (Figure 4D), while there was a non-significant trend towards higher P70S6K expression in patients with progressive disease (Figure 4E) and a later disease stage (Figure 4F). In 8 patients with matched diagnosis and relapse samples, the levels of P70S6K expression were also significantly increased upon relapse (Figure 4G), further highlighting the functional role of P70S6K in the cellular response to platinum therapy.

### P70S6K promotes cell cycle arrest in response to cisplatin

As P70S6K has a known role in cell cycle progression (Lane et al., 1993), we performed live cell imaging of the FUCCI two-colour sensor of cell cycle progression (Sakaue-Sawano et al., 2008) across the resistant A549, NCI-H1573 and NCI-H358 cell lines. This approach allowed us to track the cell cycle progression and fate of individual cells for 72 h following treatment with a 2 h pulse of cisplatin (Figure 5A). In this assay, treatment with a single pulse of cisplatin caused several notable cell cycle responses. First, A549 (*TP53* wildtype) and NCI-H358 (*TP53* null) cells were significantly more likely to remain arrested in G1, both before (G1 arrest before mitosis; ABM) and after undergoing mitosis (G1 arrest after mitosis; AAM) (Figure 5B,C). In contrast, NCI-H1573 cells did not show any significant increase in G1 transit time, as may be expected from a *TP53* mutant cell line.

**Figure 5:**
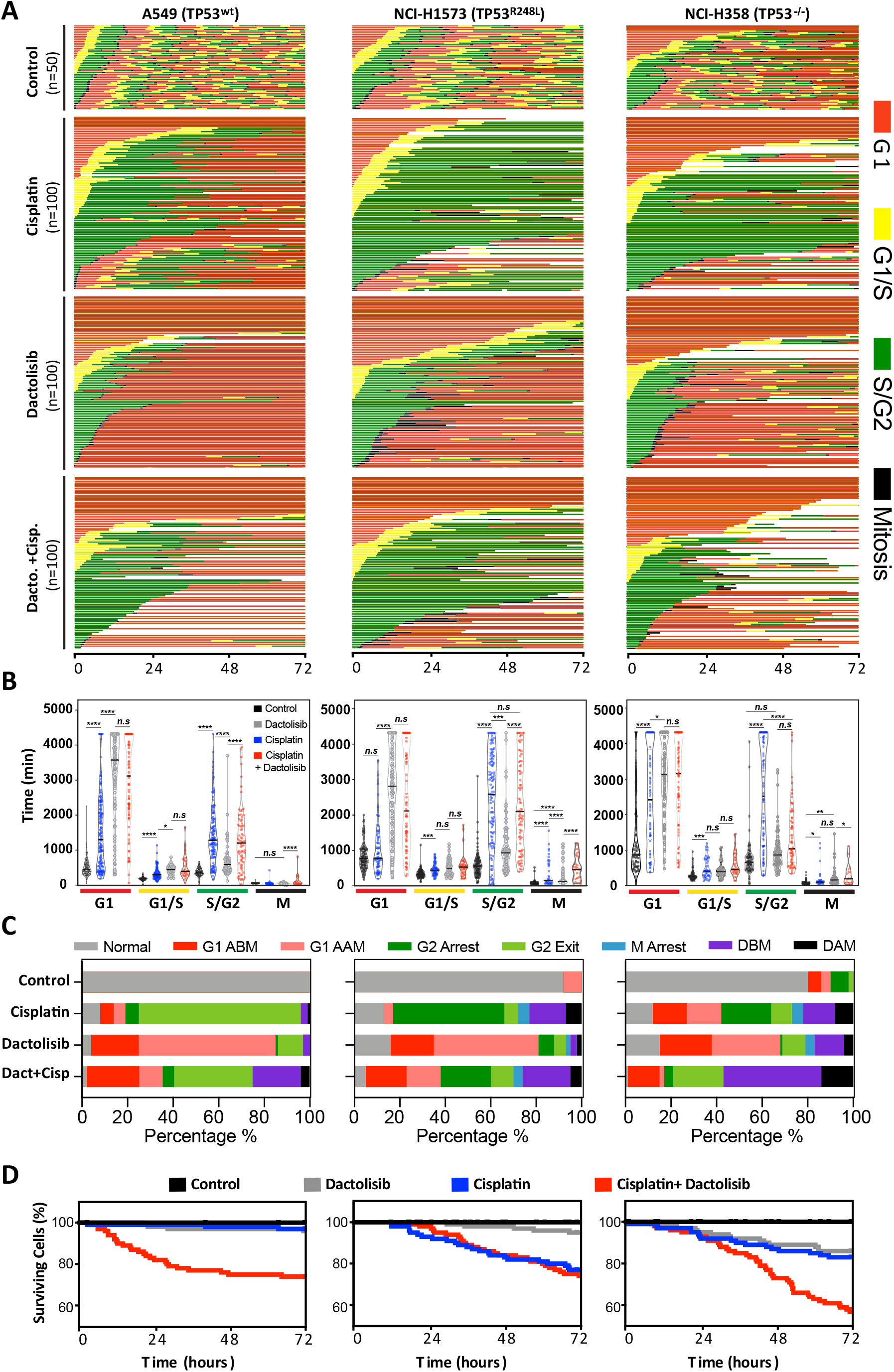
FUCCI analysis of cell cycle progression. **(A)** Live-cell imaging of the FUCCI biosensor proteins mVenus-hGeminin(1/110) and mCherry-hCdt1(30/120), stably expressed by the A549, NCI-H1573 and NCI-H358 cell lines. Images were taken every 20 min for 72 h under control conditions, or following a cisplatin pulse (5 µg/mL, 2 h) in the presence or absence of dactolisib (1 µM). **(B)** Quantification of the length of each cell cycle phase under each treatment condition (n=17-175, mean ± SD), **** p<0.0001, *** p<0.001, ** p<0.01, * p<0.05. (**C**) Quantification of fate of each cell; including G1 arrest before mitosis (G1 ABM), G1 arrest after mitosis (G1 AAM), death before mitosis (DBM) and death after mitosis (DAM). (D) Survival curves indicating the proportion of viable cells over time under each treatment condition.

For all cell lines, cisplatin pulsing resulted in a significant increase in S/G2 transit time (Figure 5B,C). Notably, the NCI-H1573 (*TP53* mutant) and NCI-H358 (*TP53* null) cells delayed for more time in G2 (G2 arrest; mean 2443 min and 2555 min respectively) compared to A549 cells (mean 1426 min), after which the mutant and null cells often entered a prolonged aberrant mitosis (Supplementary Figure 5), resulting in a small increase in cells dying during mitosis or in the following G1/S phase (Figure 5C, death after mitosis; DAM). In contrast, A549 cells rarely entered into mitosis, instead many of these S/G2 arrested cells turned from green (S/G2) back to red (G1) without undergoing mitosis (G2-exit) (Supplementary Figure 5). A similar G2-exit, senescence state has been reported to be dependent on p21 (Gire & Dulic, 2015) and likely provides *TP53* wildtype cells protection from death by preventing progression through an aberrant mitosis.

Treatment of all cell lines with dactolisib as a single agent also caused a significant increase in G1 arrest both before (G1 ABM) and after mitosis (G1 AAM), irrespective of *TP53* status (Figure 5B,C). Consequently, for all cell lines, dactolisib treatment of cells within early G1 phase during the cisplatin pulse resulted in many cells remaining arrested and viable in G1 (G1 ABM), highlighting the importance of DNA replication for inducing cisplatin induced toxicity and killing. However, for the *TP53* wildtype and null cells that were in late G1 or S phase at the time of cisplatin pulsing, dactolisib treatment significantly reduced S/G2 transit time (Figure 5B) and greatly increased the number of cells that underwent apoptosis before mitosis (Death before mitosis; DBM) (Figure 5C,D). Notably, dactolisib treatment did not prevent cisplatin treated cells from entering into an aberrant mitosis, but did significantly increase the duration of mitotic arrest, independent of *TP53* status (Figure 5B), which correlated with an increase in death during or after mitosis (DAM) in *TP53* wildtype and null, but not mutant cells (Figure 5C,D). Surprisingly, dactolisib treatment did not result in any sensitisation of *TP53* mutant H1573 cells to cisplatin (Figure 5D), which also corresponded with a failure of dactolisib to significantly reduce S/G2 phase transit time (Figure 5B).

In summary, combination therapy with dactolisib sensitised actively cycling cells to cisplatin through distinct mechanisms dependent on *TP53* status. In the *TP53* wildtype A549 cells it induced pre-mitotic cell death and prevented cells from undergoing a protective G2-exit, which is likely partially dependent on the p53-p21 axis. Similarly, in the *TP53* null line, dactolisib promoted both pre-mitotic cell death and the entry of cells into a prolonged deleterious aberrant mitosis, the latter likely due to a weakened p53-dependent G2 checkpoint(Engeland, 2018). While in *TP53* mutant H1573 cells, there was no significant sensitisation due to a sustained S/G2 arrest in the presence of dactolisib.

### Response to dactolisib is dependent upon TP53 status

A notable observation arising from this cell cycle analysis was the ability of dactolisib to increase apoptosis by preventing an unexpected cell cycle arrest in the *TP53* null NCI-H358 cell line. Interestingly, this association between cell cycle arrest and cisplatin resistance can also be observed from the original analysis of signalling dynamics (Figure 1G, Supplementary Figure 3). While both the NCI-H1299 and NCI-H358 cell lines are *TP53* null, there was an increased expression of p21 following cisplatin treatment in the resistant NCI-H358 line, but not the sensitive NCI-H1299 line. Importantly, the increased p21 expression by NCI-H358 cells was also observed by western blotting of independent samples and was efficiently inhibited by both treatment with dactolisib (Figure 6A,B) and the specific knockdown of P70S6K with siRNA (Figure 6C). Taken together, this data suggests that P70S6K activity is necessary for promoting p21 expression in order to maintain cell cycle arrest following cisplatin treatment. However, in the absence of p53, we observed that this can be mediated by the related transcription factor p63, which is elevated in the NCI-H358 line following treatment with cisplatin, and also inhibited by the addition of dactolisib (Figure 6A,B) or treatment with P70S6K siRNA (Figure 6C).

**Figure 6:**
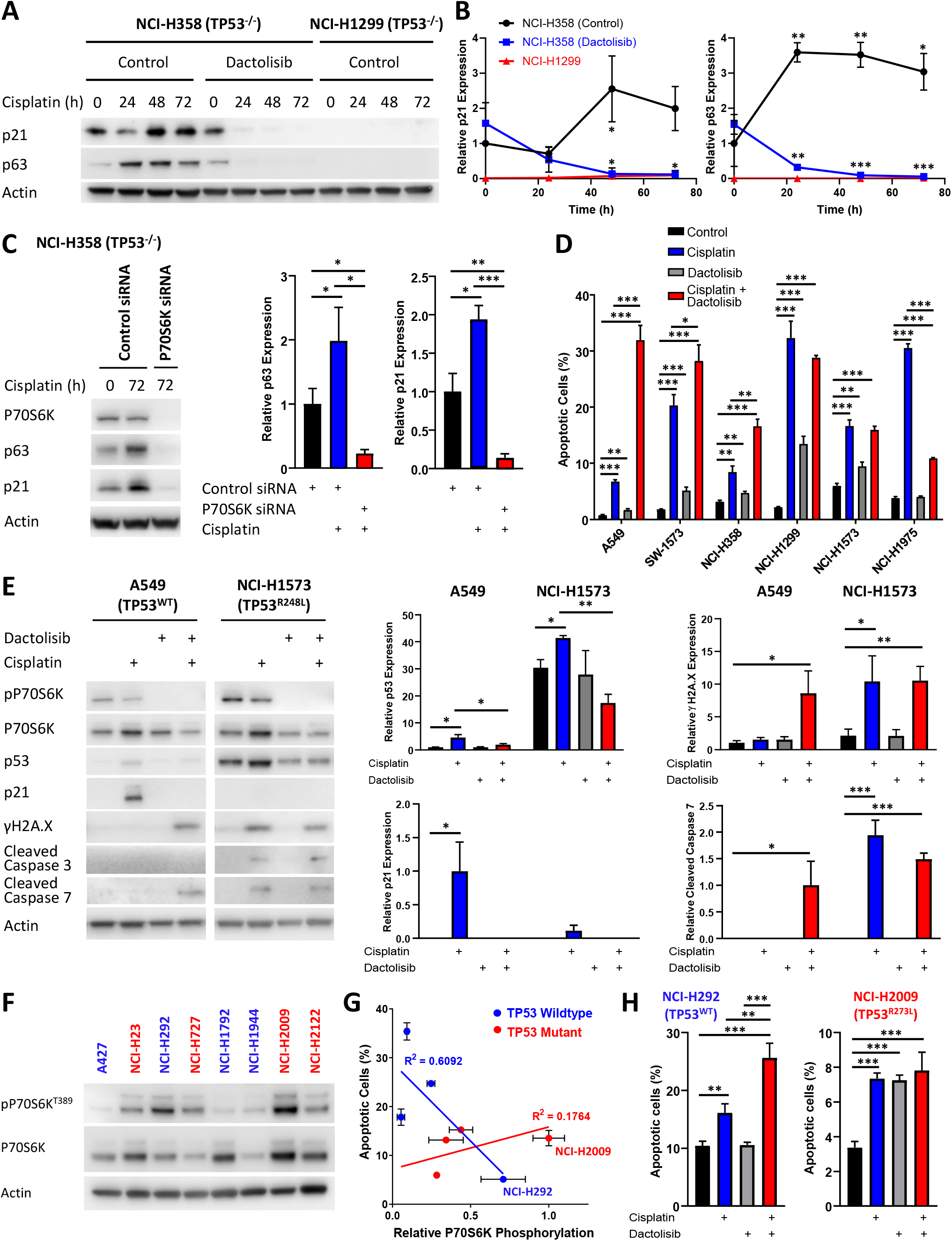
Response to dactolisib is dependent upon *TP53* mutation status. **(A)** Western blotting on lysates from NCI-H358 and NCI-H1299 cells following a cisplatin pulse, with the addition of dactolisib (1 µM), as indicated. **(B)** Quantification of effects of dactolisib upon cisplatin induced p63 and p21 expression (n=3, mean ± SD). **(C)** Western blotting on lysates from NCI-H358 cells, treated with P70S6K or control siRNA, as indicated, prior to and following a cisplatin pulse (n=3, mean ± SD). **(D)** Apoptosis measured by propidium iodide staining for the sub-G1 population performed 72 h following a cisplatin pulse with the addition of dactolisib (1 µM) as indicated (n=3, mean ± SD). **(E)** Western blotting on lysates from A549 and NCI-H1573 cells, 48 h following a cisplatin pulse, with the addition of dactolisib (1 µM), as indicated (n=3, mean ± SD). **(F)** Western blotting across a second panel of lung adenocarcinoma cell lines. **(G)** Correlation between P70S6K phosphorylation and apoptosis measured by propidium iodide staining for the sub-G1 population, performed 72 h following a cisplatin pulse (n=3, mean ± SD).

Given the potential role for p21 in mediating resistance to cisplatin in the NCI-H358 line, we investigated whether sensitisation by dactolisib would be generalizable across the original panel of 6 lung adenocarcinoma cell lines with differing *TP53* status. In accordance with this hypothesis, and in line with the FUCCI single-cell imaging (Figure 5A), both the *TP53* wildtype cell lines were significantly sensitised to cisplatin by the addition dactolisib, as was the resistant *TP53* null line, NCI-H358 (Figure 2D, 2H, 6D). The sensitive *TP53* null line, NCI-H1299, was not further sensitised and appeared to already be at the upper limit of apoptosis within this model. Also, in line with the FUCCI analysis, dactolisib was not able to increase cisplatin-induced apoptosis in the *TP53* mutant line NCI-H1573, whilst it significantly antagonised cisplatin-induced apoptosis in the *TP53* mutant NCI-H1975 line.

Lending further support to the key role of p21 in mediating cell cycle arrest, and thereby preventing DNA damage induced apoptosis, the addition of dactolisib inhibited p53 and p21 accumulation in the *TP53* wildtype A549 cell line, resulting in elevated γH2A.X expression and caspase 3 cleavage following cisplatin treatment (Figure 6E). However, in the *TP53* mutant NCI-H1573 line, which was not sensitised by dactolisib, p21 expression was not significantly elevated following cisplatin treatment, meaning that γH2A.X expression and caspase 3 cleavage were already elevated and could not be inhibited by dactolisib (Figure 7).

**Figure 7:**
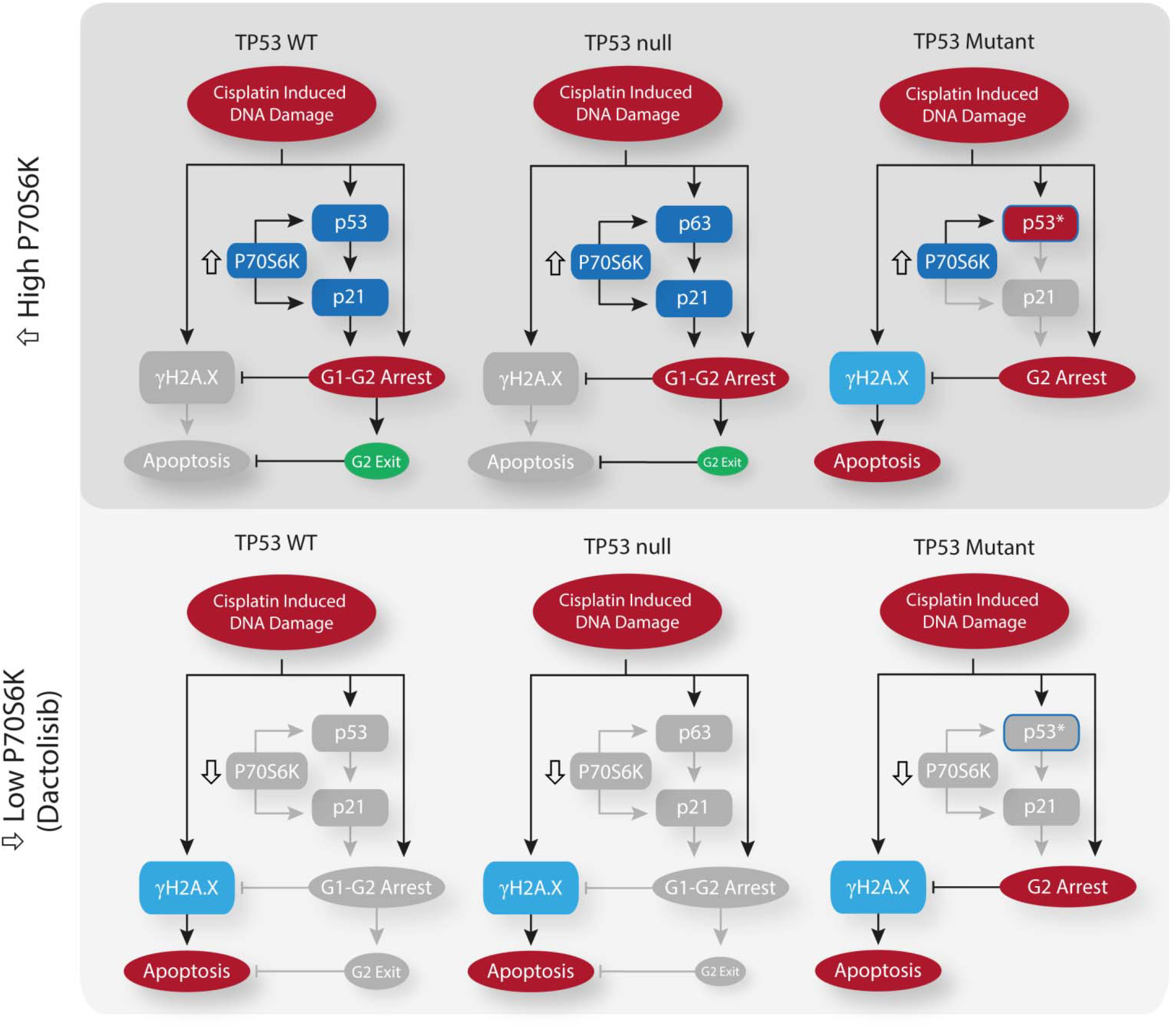
Schematic outlining the effect of P70S6K on cisplatin induced cell cycle arrest on *TP53* mutation specific backgrounds. Schematic outlining the effect of cisplatin on cell cycle arrest, DNA damage and apoptosis across the spectrum of *TP53* mutation states, and the influence of P70S6K expression levels or inhibition upon these processes.

To further validate this link between sensitisation by dactolisib and *TP53* mutation status, we took another panel of 8 lung adenocarcinoma cell lines (Supplementary Table 4) and correlated their relative expression of phosphorylated P70S6K (Figure 6F) to their apoptotic response to cisplatin (Figure 6G). This secondary analysis revealed a general trend towards higher levels of phosphorylated P70S6K and decreased apoptosis in response to cisplatin across the whole panel. However, the strength of this correlation was greatly increased when considering the *TP53* status of the lines, with a strong correlation observed for wildtype *TP53* lines (r^2^ = 0.6092) but no correlation for *TP53* mutant lines (r^2^ = 0.1764). In agreement with our finding that elevated P70S6K activity promotes resistance in a *TP53* dependent manner, combination therapy with cisplatin and dactolisib significantly sensitised the high pP70S6K/*TP53* wildtype NCI-H292 cell line, but not the high pP70S6K/*TP53* mutant NCI-H2009 line. Therefore, this data demonstrates that P70S6K promotes p21-dependent cell cycle arrest following cisplatin treatment, presumably through enhanced translation of p53/p21 in *TP53* wildtype cell lines and p63/p21 in *TP53* null lines (Figure 7). However, in the presence of mutant p53, P70S6K is not able to influence this DNA damage induced signalling axis and is therefore not associated with sensitivity to cisplatin.

## Discussion

A number of studies have been performed investigating the potential mechanisms of resistance to platinum-based chemotherapies (Stewart, 2007), although this has yet to result in the identification of clinically successful combination therapies for lung adenocarcinoma. One potential explanation for the high number of *in vitro* findings that have not translated to the clinic is the use of experimental techniques that do not replicate the *in vivo* pharmacokinetics of platinum therapies (Shen et al., 2012). To date, most *in vitro* approaches have involved continuously culturing cell lines in the presence of high doses of platinum chemotherapy for multiple days, potentially allowing for more extensive DNA damage and off-target effects that would not be seen *in vivo*. Here we have utilised a pulse model that more accurately models both the concentration and timing of cisplatin that would be observed clinically. This is an important consideration, as platinum therapies are known to act via the formation of DNA adducts, although they are also capable of bonding with proteins and RNA (Jamieson & Lippard, 1999). It is likely that the continuous culturing of cells in the presence of high doses of these drugs *in vitro* would result in the accumulation of numerous off-target adducts and the activation of a stress response incongruent with the actual *in vivo* apoptotic mechanism. Indeed, this effect may be apparent in Figure 1C, where p38 activity is induced by the continuous exposure model, but not by the pulse model. A number of previous studies have implicated the activity of p38 in mediating resistance to cisplatin (Galan-Moya et al., 2011, Hernandez Losa et al., 2003, Pereira et al., 2013, Sarin et al., 2018), however we did not observe highly elevated p38 signalling in any cell line following the cisplatin pulse (Figure 1F), nor a correlation between p38 activity and the apoptotic response (Figure 2).

Instead, through the use of this pulse model, and by overlaying real-time apoptosis data onto a multivariate signalling analysis across a panel of lung adenocarcinoma lines, we have now identified elevated P70S6K activity as an effector of inherent platinum chemoresistance. High levels of P70S6K expression have previously been associated with aggressive tumour behaviour in lung cancer (Chen et al., 2017), as well as other cancer types such as breast (Holz, 2012), colorectal (Nozawa et al., 2007) and liver (Sahin et al., 2004) cancers. This is also supported by our analysis of P70S6K expression patient samples (Figure 4), where high mRNA and protein levels were both associated with poor overall survival, and P70S6K protein expression was significantly elevated in relapsed tumours. Importantly, we also propose that combination therapy with the dual PI3K/mTOR inhibitor dactolisib can sensitise lung adenocarcinomas to platinum chemotherapy, although only in the context *TP53* WT or *TP53* null tumours. This detailed mechanistic understanding is especially important given that P70S6K amplification can occur on a background of either mutant or wildtype *TP53* (Figure 4A).

Functionally, dactolisib will result in the inhibition of a number of substrates downstream of PI3K and mTOR. However, as the specific knockdown of P70S6K with siRNA recapitulated the effects of dactolisib on p63, p21, γH2A.X, cleaved caspase 3 and apoptosis, it is likely that the elevated expression and activity of P70S6K in resistant lung adenocarcinoma cells is the functional target of dactolisib and a causal effector of platinum resistance in these cells. Further supporting this central role of P70S6K, previous research has demonstrated that P70S6K can regulate cell cycle progression via the enhanced translation of p21 mRNA (Guegan et al., 2014), although our data also suggest that P70S6K may also play a role in the enhanced expression of both p53 and p63 in this context.

The combination of cisplatin and dactolisib has also previously been proposed for osteosarcoma (Huang et al., 2018), head and neck squamous cell carcinoma (Hsu et al., 2018) and bladder cancer (Moon du et al., 2014). However, our finding that this combination therapy is effective in *TP53* WT and null, but not *TP53* mutant tumours, has significant implications for the clinical application of these drugs and the design of potential clinical trials. The failure of many clinical trials is frequently attributed to the lack of suitable biomarkers to stratify the patient cohort (Hay et al., 2014), highlighting the value of our approach that is capable of dissecting the mechanism of variable drug response across differing genetic backgrounds. The inability of dactolisib to regulate p21 expression in *TP53* mutant tumour cells following cisplatin treatment likely underlies the inability of dactolisib to sensitise these cells to cisplatin. Therefore, within the context of a potential clinical trial for this drug combination, the *TP53* mutation status of each patient tumour would be a central consideration for identification of the most effective treatment strategy and an understanding of individual patient response. However, this finding is also widely applicable to the understanding of how cancer-related signalling networks govern drug response and chemoresistance, and how a thorough characterisation of these dynamics will be necessary for the proper design and implementation of any consistently effective, patient-specific therapy.

## Materials and Methods

### Antibodies, Plasmids, and Reagents

The cisplatin modified DNA (ab103261) and γH2A.X (ab26350) antibodies were from Abcam (Cambridge, USA). The phospho-STAT3 S727 (#9134), phospho-Akt S473 (#9271), phospho-NF-κB S536 (#3033), pERK T202/Y204 (#9101), pATF2 T71 (#5112), (P70S6K (#2708), phospho-P70S6K T389 (#9205), p21 (#2947), cleaved caspase 3 (#9661) and cleaved caspase 7 (#9491) antibodies were from Cell Signaling (MA, United States). The γH2A.X (S139) antibody (AB26350) was from Abcam (MA, USA). The p53 antibody (sc-126) was from Santa Cruz Biotechnology (TX, USA). The p63 antibody (NB100-691) was from Novus Biologicals (CO, USA). The actin monoclonal antibody (AC-15) was from Sigma-Aldrich (MO, United States). The SignalSilence p70/85 S6 Kinase siRNA was from Cell Signaling (MA, United States). The plasmids for FUCCI live cell imaging, mVenus-hGeminin(1/110) and mCherry-hCdt1(30/120), were a kind gift from Dr Atsushi Miyawaki (Riken, Japan). Dactolisib (NZP-BEZ235), MK2206, S3I-201, UO126 and SC75741 were all from Selleck Chem (MA, USA).

### Cell lines

All lung adenocarcinoma cell lines have been previously described (Marini et al., 2018). The lines were cultured in Advanced RPMI (serial no) containing 1% FCS and 1% GlutaMAX (serial no) under standard tissue culture conditions (5% CO2, 20% O2). All cell lines were authenticated by short tandem repeat polymorphism, single-nucleotide polymorphism, and fingerprint analyses, passaged for less than 6 months.

Stable cell lines expressing the FUCCI biosensor were generated as previously described (Sakaue-Sawano et al., 2008) using mVenus-hGeminin(1/110) and mCherry-hCdt1(30/120) probes. Briefly, cells were first transduced with mVenus-hGeminin(1/110) lentiviral particles. Cells were FACS sorted based upon Venus fluorescence, and the resulting cell population transduced again with mCherry-hCdt1(30/120) lentiviral particles. mCherry positive cells were FACS sorted, resulting in a double positive population used for live cell imaging.

### Flow cytometry

Samples for flow cytometry were fixed in −20°C ethanol overnight, and then resuspended in a DNA staining solution containing 10mg/mL RNaseA and 1mg/mL propidium iodide for 30 min before analysis. Flow cytometry was performed using BD FACS Canto II system.

### Confocal Microscopy

For the visualisation of cisplatin-DNA adducts following treatment with a pulse of cisplatin (5 μg/mL, 2 h), cells were grown on Histogrip (Life Technologies) coated glass coverslips. Prior to fixation, cells were treated with 1 % Triton X-100 for 1 min, fixed with ice-cold 100% Methanol and stored overnight at −20°C. The cells were then permeabilized with 0.5% Triton X-100 for 10 min, washed twice with PBS and incubated for 30 min with 2M HCl to denature the DNA. Cells were washed again and incubated in 1% H_2_O_2_ to quench the endogenous peroxidases, before blocking for 30 min and incubated with primary antibody (1:500) overnight at 4 C in blocking solution. The following day, cells were incubated with Secondary-Biotinylated Ab at RT, before incubating them with ABC solution (Vectastain elite ABC HRP kit, Vector Laboratories). Cells were later incubated with TSA solution (TSA plus Cyanine 3 System, PerkinElmer). DNA was stained with H33342 and images collected using a Leica DMI5500 (40x magnification). Images were quantified using Fiji Software. Briefly, color images were split into separate channels. H33342 channel was used to identify the nucleus and generate a mask that was placed on top of the CisPt-DNA channel to quantify the signal coming from the nuclear area. Cytoplasmic area was manually identified in each cell and signal quantified. Background signal was obtained and subtracted from the nuclear and cytoplasm signal. From this data, a nuclear/cytoplasmic ratio was obtained for between 100 and 200 cells at each time point.

### MAGPIX multiplex assays

Multiplex analysis was performed using a Bio-Plex MAGPIX system (#171015044) and Bio-Plex Pro-Wash Station (Biorad). All cell lysates were prepared using standard cell lysis buffer (50mM Tris HCl pH7.4, 150mM NaCl, 1mM EDTA, 1% (v/v) TritonX-100) supplemented with protease and phosphatase inhibitors. Lysates were analysed on all kits, according to manufacturer’s instructions. Data was generated using Bio-Plex Manager MP and analysed on the Bio-Plex Manager 6.1 software.

Lysates were analysed using the Milliplex map DNA Damage/Genotoxicity Magnetic Bead Panel (7-plex), Milliplex map Early Apoptosis Magnetic Bead (7-plex), Milliplex map TGFβ Signalling Pathway Magnetic Bead kit (6-plex), Bio-Plex Pro Cell Signalling Akt Panel (8-plex), Bio-Plex Pro Cell Signalling MAPK Panel (9-plex), Bio-Plex Pro RBM Apoptosis Panel 2 and Bio-Plex Pro RBM Apoptosis Panel 3. Individual beads were also used to analyse NF-κB (Ser536), IκBα (Ser32/36), c-Jun (Ser63), total P53 and cleaved PARP (Biorad). The data was normalized to the median value at the 0 h time point for each analyte and a log transformation was conducted on the resulting dataset. The principal component analysis was performed using MATLAB and Statistics Toolbox Release 2019a (The Mathworks, Inc., Natick, Massachusetts, United States of America)

### Live-cell imaging of apoptosis

Live-cell imaging of apoptosis was performed using an Essen Bioscience IncuCyte ZOOM Live-Cell Analysis System and a Thermo Fisher Scientific HERAcell 240i CO_2_ Incubator. Cells were seeded into Corning Costar TC-treated 96-Well Plates and imaged over a 72 hr period at 2 hr intervals over 4 fields of view per well. Caspase activation was visualised using 1µM NucView 488 Caspase-3 Enzyme Substrate (Biotium). Cell viability was quantified using propidium iodide (1 μg/mL). The generated images were analysed using IncuCyte ZOOM Software Version 2016B.

### Western blotting

Lysates for western blotting were prepared using standard lysis buffer (50mM Tris HCl pH7.4, 150mM NaCl, 1mM EDTA, 1% (v/v) Triton X-100) containing protease inhibitor (p8340, Sigma) and 0.2 mM sodium orthovanadate. The NuPAGE SDS PAGE Gel System and NuPAGE Bis Tris Precast Gels (4-12% and 12%) (Life Technologies) were used to perform gel electrophoresis. Western Lightning PLUS Enhanced Chemiluminescent Substrate (PerkinElmer) was used for imaging western blots on the Vilber Lourmat Fusion chemiluminescent imaging system. Quantitative western blotting was performed using multistrip western blotting, as performed previously (Kennedy et al., 2019).

### NCI-H358 xenograft model

NCI-H358 cells (2×10^6^) resuspended in 100 μL PBS:Matrigel were injected subcutaneously into the flanks of nude mice. Tumour growth was assessed twice weekly by calliper measurement and mice were randomized to treatment arms when tumours reached 150 mm3 (using the formula: width2 x length x 0.5). Carboplatin (60 mg/kg) was delivered by a single tail-vein injection. Dactolisib was prepared in 10% DMSO:90% PEG300 and administered twice-weekly at 45 mg/kg by oral gavage. All in vivo experiments, procedures and endpoints were approved by the Garvan Institute of Medical Research Animal Ethics Committee.

### Immunohistochemistry

Immunohistochemistry was performed on formalin fixed paraffin embedded sections using the Leica BOND RX (Leica, Wetzlar, Germany). Slides were first dewaxed and rehydrated, followed by heat induced antigen retrieval performed with Epitope Retrieval Solution 1 BOND (Leica, Wetzlar, Germany). Primary antibodies were diluted 1:600 (P70S6K) and 1:500 (Ki67) in Leica antibody diluent and incubated for 60 min on slides. Antibody staining was completed using the Bond Polymer Refine IHC protocol and reagents (Leica, Wetzlar, Germany). Slides were counterstained on the Leica Autostainer XL (Leica, Wetzlar, Germany). Leica CV5030 Glass Coverslipper (Leica, Wetzlar, Germany) and brightfield images were taken on the Aperio CS2 Slide Scanner (Leica, Wetzlar, Germany). Quantification of Ki67 staining was performed on three fields of view for each tumour section, and quantified using the particle analysis function of Image J (v1.49).

### FUCCI live-cell imaging

For FUCCI live cell imaging, cells were seeded on 12 well plates and imaged using a Leica DMI6000 using a 20x NA 0.4 objective. Images were taken every 20 min for 72 h. Before adding any drug (cisplatin, dactolisib or both) an image was taken in order to annotate the cell cycle phase before commencing treatment. Individual cells were tracked manually, with the colour of the nucleus annotated at each time point (Red=G1; Yellow=G1/S, Green=S/G2), the cells were also scored for nuclear envelope breakdown (NEBD) and early signs of anaphase. Mitotic length was calculated by the time period from commencement of NEBD to anaphase. Interphase length was calculated from anaphase to the next daughter cell NEBD. Only one daughter was followed and annotated. Tracking graphs were generated using Prism 7.

## Acknowledgements

The authors would like to acknowledge Dr Atsushi Miyawaki (Riken, Japan) for provision of the mVenus-hGeminin(1/110) and mCherry-hCdt1(30/120) constructs for FUCCI imaging. DRC was a Cancer Institute NSW Future Research Fellow (2013/FRL102).

## Author Contributions

JFH conducted the experiments and wrote the paper. AGR conducted the experiments. VC, RZ and JZRH performed the animal work. YEIO and MP performed western blotting experiments. VC, ADM, AN and DD collated the patient cohort and clinical data. AB established and supervised the live-cell imaging techniques, and edited the paper. DNW conceived the study, designed experiments, supervised AGR and edited the paper. DRC conceived the study, designed experiments, supervised JFH, JZRH, YEIO and MP, and wrote the paper.

## Conflict of Interests

The authors declare no competing interests.

## The Paper Explained

### PROBLEM

Platinum chemotherapy has been the cornerstone of treatment for lung adenocarcinoma for the last four decades, however response rates still remain below 30%. Attempts to understand the mechanisms of innate resistance to these chemotherapy agents have been hampered by *in vitro* approaches that do not recapitulate *in vivo* pharmacokinetics. Therefore, while numerous mechanisms of resistance have been proposed, they have not translated to clinically viable options to overcome platinum resistance.

### RESULTS

Using a cisplatin pulse model that closely mimics the timing and concentration of *in vivo* drug exposure, we firstly demonstrated that the apoptotic and signalling response induced by this pulse model was fundamentally different to that observed when cells are cultured in the continuous presence of cisplatin. Therefore, we further utilised this pulse model to profiled cisplatin-induced signalling, DNA damage and apoptotic responses across a panel of lung adenocarcinoma cell lines. Overlaying a multi-dimensional analysis of this dataset with real-time, single-cell imaging of apoptosis and cell cycle progression facilitated the identification of a P70S6K signalling axis in resistant cells, that directly opposed a late phase apoptotic response. The inhibition of P70S6K using either siRNA, or the dual PI3K/mTOR inhibitor dactolisib, sensitised resistant cells to cisplatin by preventing cell cycle arrest and promoting further DNA damage and apoptosis. Detailed mechanistic analysis demonstrated that as this cell cycle arrest was dependent upon enhanced p21 expression, this combination therapy was effective on a *TP53* wildtype or *TP53* null background, but not in the presence of mutant *TP53*.

### IMPACT

The failure of many clinical trials is frequently attributed to the lack of suitable biomarkers to stratify the patient cohort. Here we demonstrate that the combination of cisplatin and dactolisib will effectively overcome platinum resistance in a strictly defined set of *TP53* wildtype and *TP53* null lung adenocarcinomas. This finding highlights the value of developing *in vitro* models that more closely match the physiological pharmacokinetics of drug exposure, and also the importance of a thorough mechanistic understanding of the signalling networks that govern drug response for the proper design and implementation of any consistently effective, patient-specific therapy.

---

**Supplementary Figure 1:**
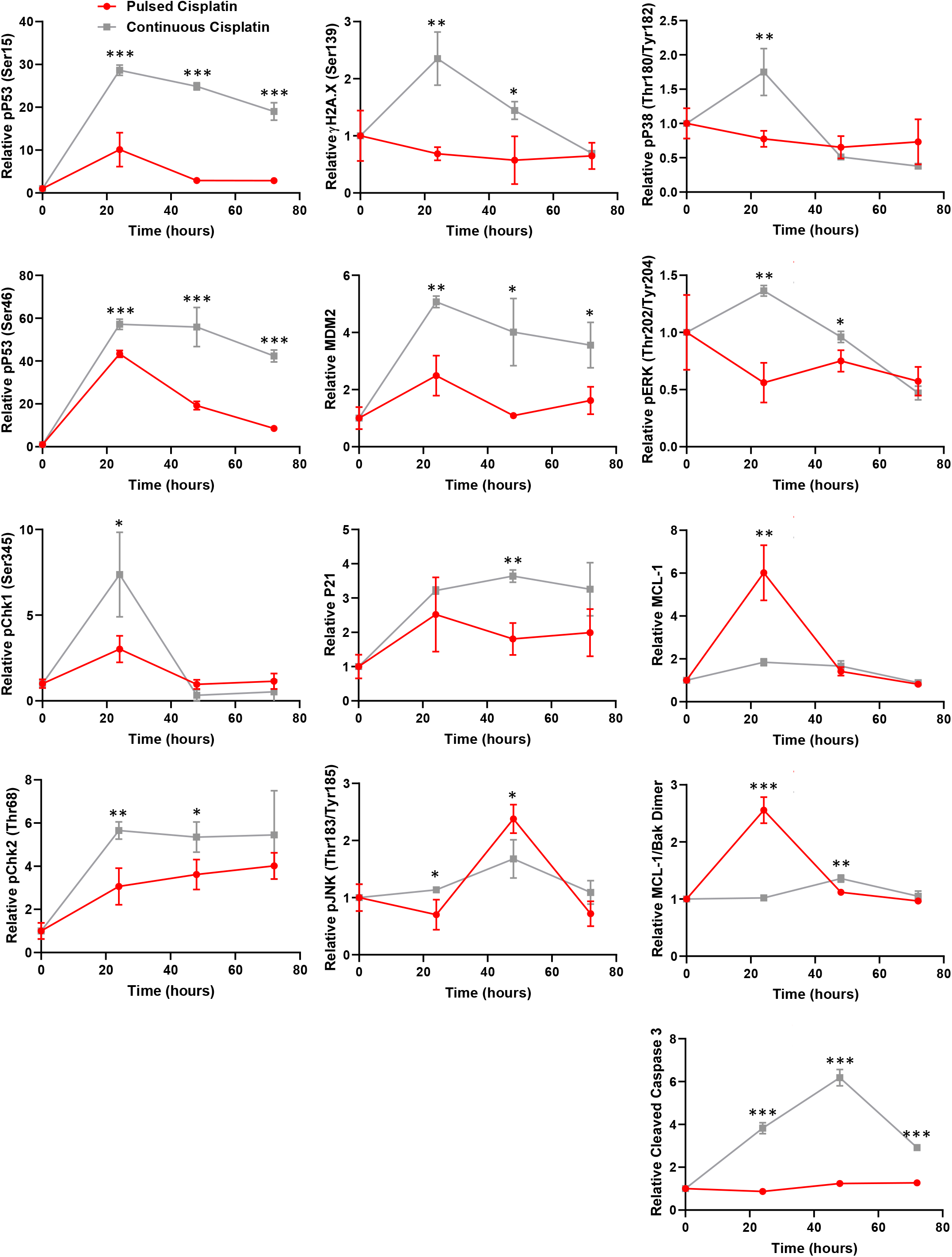
Continuous versus pulsed cisplatin treatment of A549 cells. Raw data for the timecourse, multiplex analysis of DNA damage response proteins following continuous cisplatin treatment (grey) or a cisplatin pulse (red) (5 μg/mL, 2 h) in A549 cells (n=3, mean ± SD. *** p < 0.001, ** p < 0.01, * p < 0.05).

**Supplementary Figure 2:**
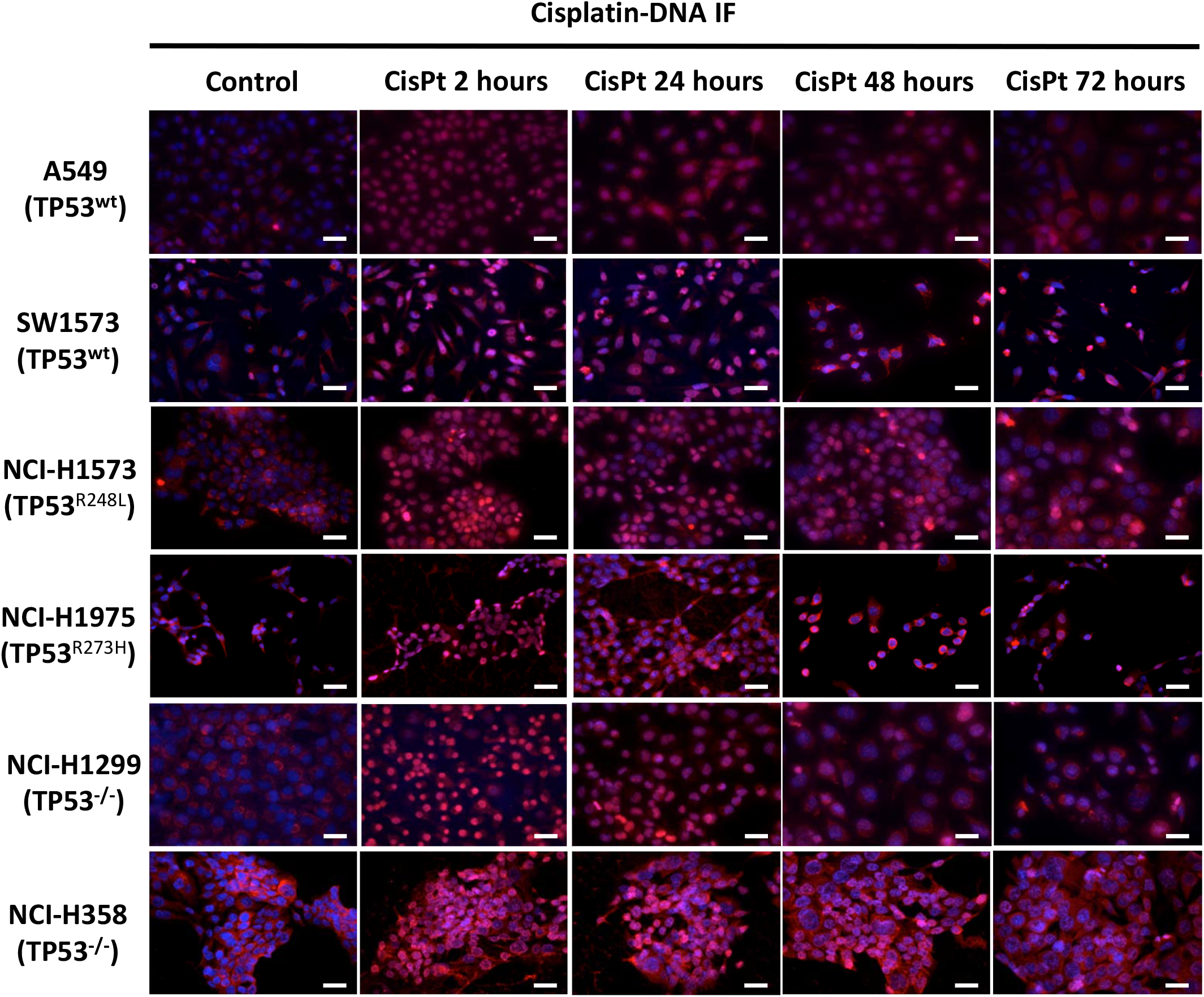
Imaging of cisplatin-DNA adducts. Representative images of anti-cisplatin antibody staining across the cell line panel following a cisplatin pulse (Scale bar: 40 μm).

**Supplementary Figure 3:**
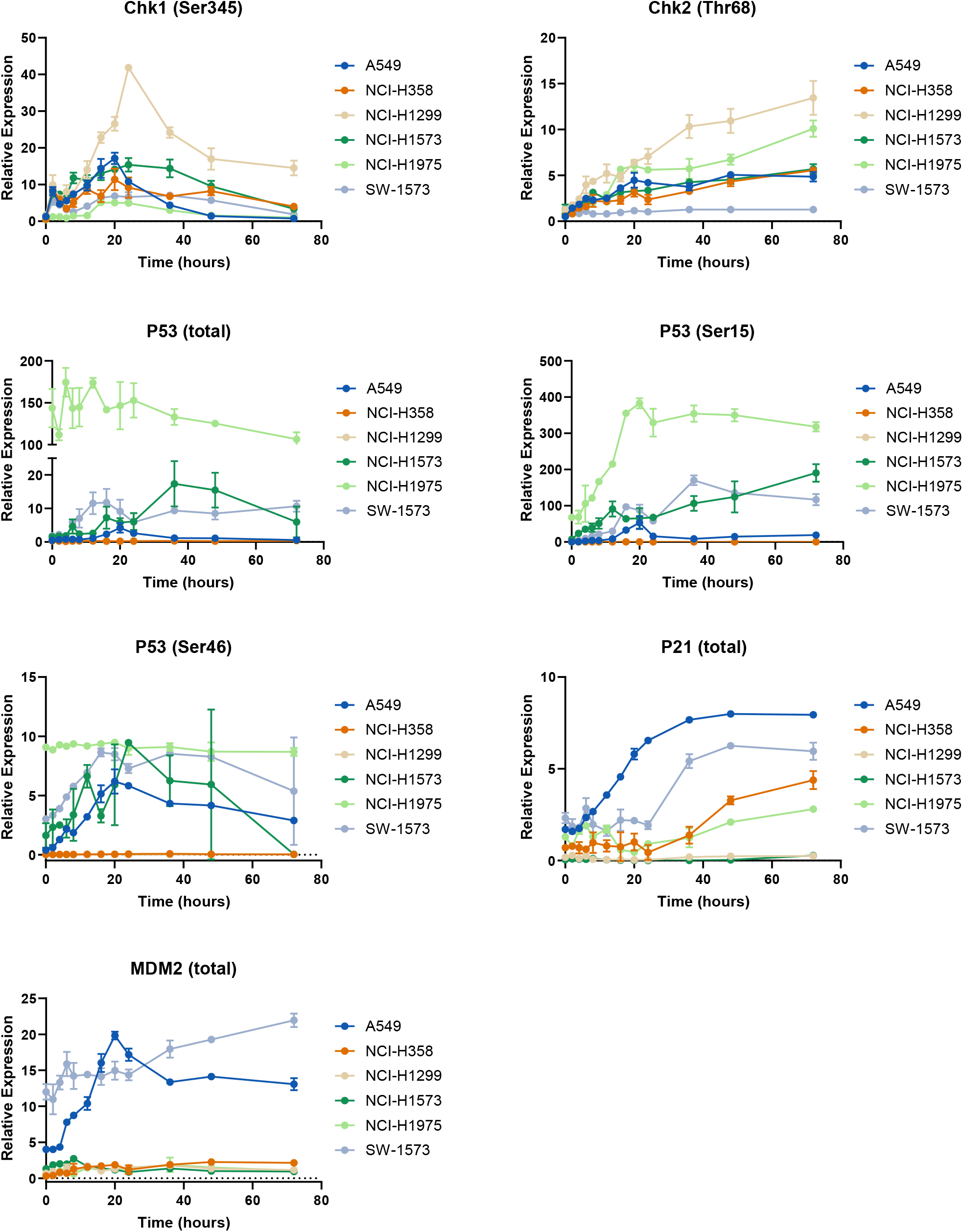
p53 pathway dynamics. Raw data for the timecourse, multiplex analysis of DNA damage response proteins following a cisplatin pulse (5 μg/mL, 2 h) across a panel of cell lines, as indicated (n=3, mean ± SD).

**Supplementary Figure 4:**
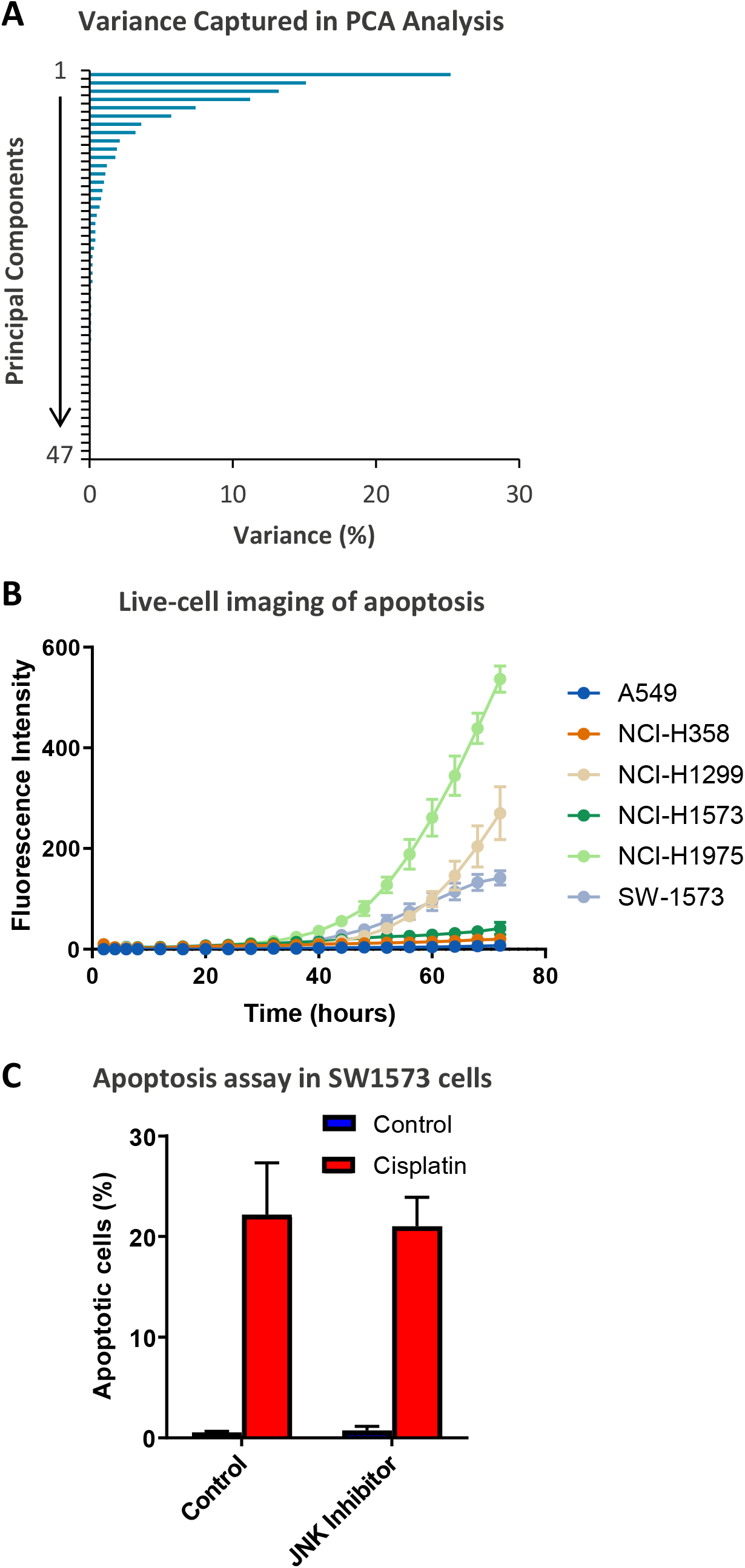
**(A)** The variance captured within each principal component of the principal component analysis (PCA), presented in Figure 2. **(B)** Live-cell imaging of apoptosis across the cell lines indicated on the Incucyte platform using a fluorescent caspase substrate (1 μM) (n=3, mean ± SD). **(C)** Apoptosis measured by propidium iodide staining for the sub-G1 population in SW1573 cells, performed 72 h following a cisplatin pulse (5 μg/mL, 2 h) with the addition of a JNK inhibitor (JNK VIII, 20 µM) as indicated (n=3, mean ± SD).

**Supplementary Figure 5:**
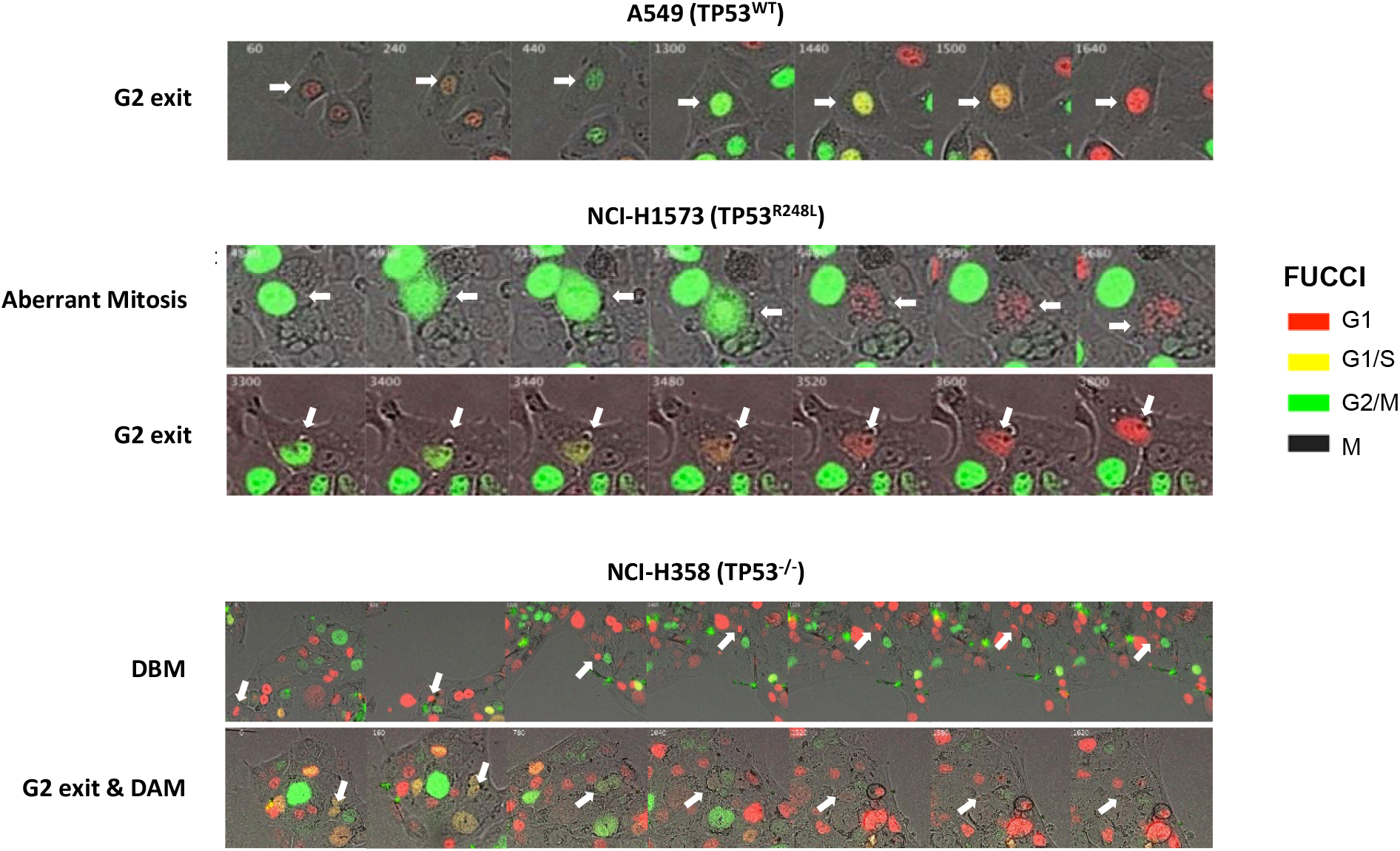
Representative images of cells expressing the FUCCI biosensor, undergoing an aberrant mitosis, G2 exit, Death before Mitosis (DBM) and Death after Mitosis (DAM) following a cisplatin pulse (5 μg/mL, 2 h).

**Supplementary Table 1:** Mutation status of the lung adenocarcinoma cell panel.

| Cell line | TP53 | RAS | EGFR | CDKN2A |
| --- | --- | --- | --- | --- |
| A549 | WT | KRAS G12S | WT | Del (Hom) |
| SW-1573 | WT | KRAS G12C | WT | Del (Hom) |
| NCI-H1573 | R248L (Hom) | KRAS G12A | WT | WT |
| NCI-H1975 | R273H (Hom) | WT | T790M,<br>L858R | WT |
| NCI-H358 | Del (Hom) | KRAS G12C | WT | WT |
| NCI-H1299 | Del (Hom) | NRAS Q61K | WT | WT |

**Supplementary Table 2:** Summary of the analytes used for multiplex signalling analysis.

| Pathway | Analytes |
| --- | --- |
| DNA Damage Response | ATR (total), Chk1 (Ser345), Chk2 (Thr68), P53 (total), P53(Ser15), P53 (Ser46), MDM2 (total), P21 (total), γH2AX (Ser139), cleaved PARP |
| AKT/mTOR Signalling | IRS-1 (Ser636/639), Akt (Ser473), BAD (Ser136), BAD (Ser112), Bcl-2 (Ser70), GSK-3$\alpha/\beta$ (Ser21/9), S6 ribosomal protein (Ser235/236), PTEN (Ser380), mTOR (Ser2448), P70S6K (Thr389) |
| MAPK Signalling | MEK1 (Ser217/221), ERK1/2 (Thr202/Tyr204), P90RSK (Ser380), P38 (Thr180/Tyr182), HSP27 (Ser78), JNK (Thr183/Tyr185), c-Jun (Ser63), ATF2 (Thr71), STAT1 (Tyr701), STAT3 (Ser727) |
| TGFβ Signalling | TGFβRII (total), Smad2 (Ser465/467), Smad3 (Ser423/425), Smad4 (total) |
| NF-κB Signalling | NF-κB (Ser536), IκBα (Ser32/36) |
| Apoptosis Regulators | BAD (total), Bcl-xL (total), BIM (total), MCL-1 (total), Bcl-xL/BAK dimer (total), MCL-1/BAK dimer (total), BAX/Bcl-2 dimer (total), Survivin (total) |
| Apoptosis Effectors | Active Caspase-8, Active Caspase-9, Active Caspase-3 |

**Supplementary Table 3:** Characteristics of the patient cohort used to generate immuno-histochemical data. CI = confidence interval.

|  |  | **n** | **%** |
| --- | --- | --- | --- |
| ***Total Samples*** |  | 52 |  |
| ***Histology*** | Adenocarcinoma | 50 | 96.2% |
|  | Large cell favouring adenocarcinoma | 1 | 1.9% |
|  | Mucinous adenocarcinoma | 1 | 1.9% |
| ***Stage*** | IV | 36 | 69.3% |
|  | IIIA | 8 | 15.4% |
|  | IIIB | 5 | 9.6% |
|  | III | 1 | 1.9% |
|  | IIB | 1 | 1.9% |
|  | II | 1 | 1.9% |
| ***Chemotherapy*** | Carboplatin/Gemcitabine | 28 | 53.8% |
|  | Carboplatin/Pemetrexed | 3 | 5.8% |
|  | Carboplatin/Pemetrexed +/- Pembrolizumab | 4 | 7.7% |
|  | Carboplatin/Taxol + Bevacizumab | 2 | 3.8% |
|  | Concurrent Carboplatin/Taxol | 11 | 21.2% |
|  | Concurrent Cisplatin/Etoposide | 3 | 5.8% |
|  | Concurrent Carboplatin + Unknown | 1 | 1.9% |

**Supplementary Table 4:**
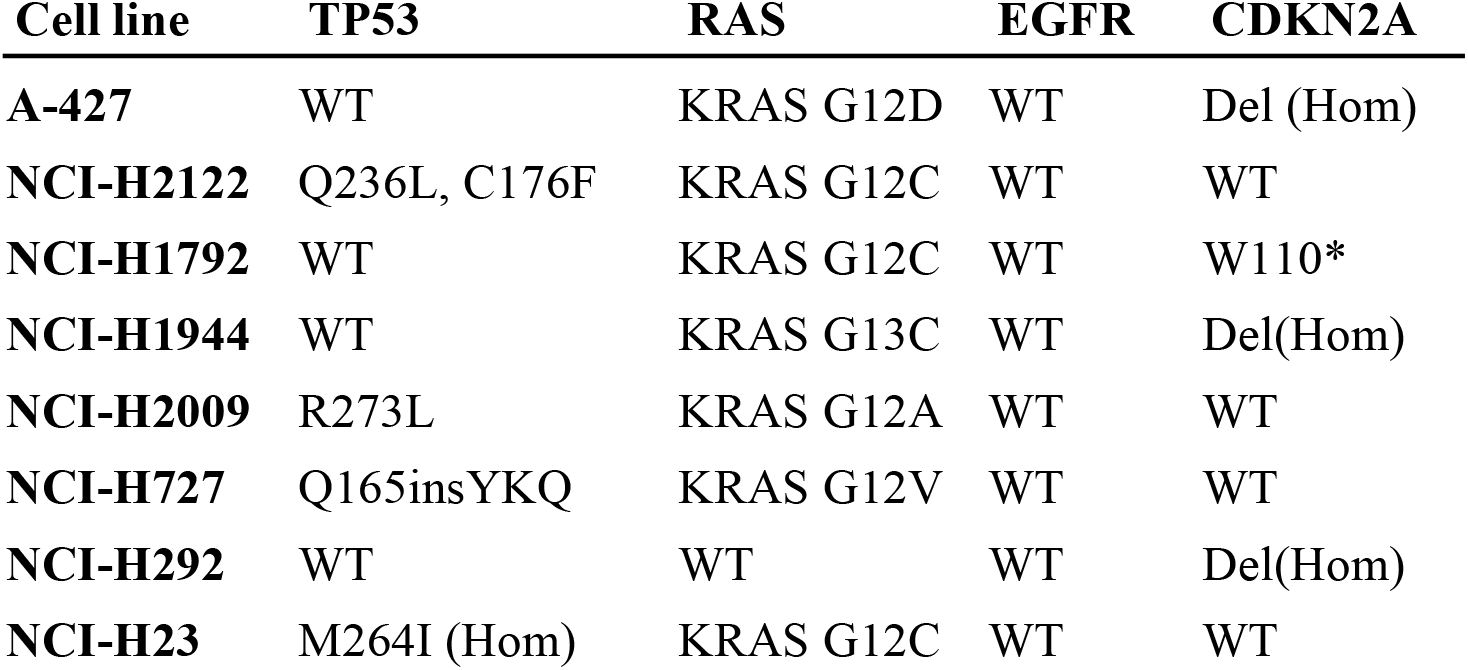
Mutation status of second lung adenocarcinoma cell panel.

